# Night-to-night sleep EEG variability over one year

**DOI:** 10.64898/2026.07.02.736125

**Authors:** Yevgenia Rosenblum, Leonore Bovy, Martin Christian Hemmsen, Jonas Duun-Henriksen, Esben Ahrens, Martin Dresler

## Abstract

This study aimed to explore night-to-night variability of multiscale sleep patterns by analyzing subcutaneous electroencephalography (sqEEG) from 20 healthy participants over one year (205–388 nights per participant, 6,429 nights in total). We utilized the time series of aperiodic slopes, sigma and slow-wave power as a new whole-night unit of sleep macrostructure. Using dynamic time warping, we calculated the distances (differences) between those time series to assess night-to-night sleep macrostructure dissimilarity. We found that the overall sleep macrostructural patterns were relatively similar across nights (20% dissimilarity), while their temporal alignment was quite variable (time series warped by ∼60% for the best alignment). Lower variation in macrostructure dissimilarity was associated with better subjective sleep quality (r=-0.25).

Then, we qualitatively compared yearlong variation in macroscale, microscale (sleep stage proportions, mean spectral power) and mesoscale (sleep cycle duration) metrics. We found that intra-individual night-to-night variation was “low” (coefficients of variation < 20%) for spectral power, sleep duration, N2 and REM sleep; “medium” (20–40%) – for N3 and macrostructure dissimilarity; and “high” (>40%) – for sleep cycle duration, wake and N1. In summary, different sleep metrics showed differential night-to-night variability, which was more metric-specific than scale-dependent. This might reflect a distinction between more trait-like versus more dynamically varying features of sleep, although this assumption needs further clarification.

**Significance statement:** The degree of the intra- and inter-individual night-to-night variability of sleep metrics at different scales shows more metric-specific than scale-dependent patterns and potentially distinguishes between trait-like and state-like features of sleep. Detailed description of the multiscale sleep structure and its night-to-night variability/stability might eventually lead to identification of electrophysiological fingerprints and bring insights about their physiological significance. The potential implications of this include the development of personalized time-sensitive recommendations for people facing (social) jet lag, shift work and other deviations from normal sleep patterns. In clinical settings, this might advance the tools to diagnose and monitor sleep alterations, predict nocturnal seizures in epilepsy care, instability in breathing control in apnea etc.

## Introduction

### Sleep EEG research

Recording polysomnography in a sleep laboratory to obtain single night snapshots has been the gold standard in basic and clinical research for decades. While this approach has shaped our current knowledge about sleep structure, it misses information on temporal dynamics and night-to-night sleep regularity, which appears to be at least as important for general and mental health as other sleep parameters, such as sleep duration (Windred et al., 2024).

Recently introduced subcutaneous electroencephalography (sqEEG) devices enable ultra-long longitudinal EEG monitoring (Duun-Henriksen et al., 2020), which has been limited thus far due to expensive costs of acquiring EEG in a professional polysomnographic laboratory and lower quality of the data acquired with EEG wearables at homes (as compared to the gold standard). Several papers describing implantable sqEEG devices (Djurhuus et al., 2023; Ahrens et al., 2024) or analyzing their data have been published thus far in patients with epilepsy (Viana et al., 2023; Helge et al., 2025) and healthy volunteers (Avigdor et al., 2025; Leguia et al., 2025). Here, we analyze sqEEG recorded in healthy participants within the Ultra-Long-Term Sleep project

(Ahrens et al., 2024), the first yearlong sleep EEG study, which as such, allows one to harness the full richness of longitudinal sleep physiology with all its nuances. This is an important advance as to date, nearly all published longitudinal sleep research is based on subjective diaries, actigraphy or different sensors (reviewed in Table S1 in Avigdor et al. (2025)), which do not include EEG and thus have no direct access to neural activity.

### Multiscale sleep structure assessment

We explore sleep structure and its variability at micro-, meso- and macroscales, assuming that different scales are interconnected. Microstructure of sleep (e.g., sleep spindles, eye movements, physiological microarousals) might dictate its mesoscale organization (NREM-REM sleep cycles), which in turn might define sleep macrostructure and vice versa. There also exist infradian (>24h) and seasonal sleep rhythmicity (Avigdor et al., 2025), but those are beyond the scope of this work.

Specifically, we first introduce a new *macroscale* metric of whole-night sleep: the EEG *time series* of aperiodic slopes as well as sigma band power and slow-wave activity (SWA) for reference. The rationale for choosing these metrics was as follows. EEG power quantifies the magnitude of the brain’s electrical activity. It consists of oscillatory and aperiodic components. The oscillatory component is subdivided into different frequency bands (e.g., slow waves, alpha, beta, sigma) associated with different cognitive states. The aperiodic component reflects non-oscillatory neural background activity. Its slope indicates how strongly low-frequency activity dominates over high-frequency activity (Bódizs et al., 2024). Aperiodic slopes are reliable markers of sleep stages and depth (Supplemental Fig.1), while time series of aperiodic slopes dynamically reflect sleep cycles (Rosenblum et al., 2025).

Second, we introduce a new measure of night-to-night macrostructure dissimilarity: the distances between successive pairs of time series assessed with *dynamic time warping* (DTW). Third, we assess variability in macrostructural dissimilarity across one year and compare it to that of commonly used sleep *micro-* and *mesoscale* metrics.

We hypothesized that night-to-night variation in sleep metrics increases with their scale, since microscale metrics are more strongly averaged (except for WASO and N1 that are averaged over few epochs), whereas macroscale metrics preserve full-night temporal dynamics, and therefore, might be quite variable and only moderately predictable.

To summarize, first, we introduce new metrics of whole-night sleep and macrostructure dissimilarity between nights; second, we qualitatively compare variability/stability of sleep metrics at different scales.

## Methods

### Dataset

We retrospectively analyzed the data collected within the Ultra-Long-Term Sleep project with implantable two-channel subcutaneous EEG (sqEEG) across one year in 20 healthy participants (Table 1). The study protocol and implantation procedure are described in the original papers (Ahrens et al., 2024; Djurhuus et al., 2023). All participants signed informed consent before the study activities. The local ethics committee approved the study, and the study was registered at clinicaltrials.gov (identifier: NCT04513743) in Denmark.

**Table 1.** Ultra-Long-Term Sleep project description.

|  |  |
| --- | --- |
| **Dataset** | Ultra-Long-Term Sleep project |
| **Reference to original papers** | Ahrens et al. (2024)<br>Djurhuus et al. (2023) |
| **No. participants** | 20 (11 females) |
| **Age, mean ± SD, median, range, years** | Mean: $32 \pm 13$, median: 28, range: 19 – 62 |
| **No. analyzed nights per participant, range** | $322 \pm 45$, range: 205 – 388 |
| **Total number of the analyzed nights** | 6,429 |
| **Device** | Subcutaneously EEG implant (24/7 EEG SubQ, UNEEG Medical A/S, Lillørød, Denmark) |
| **Electrode** | An electrode with three contacts: TP9/TP10, T9/T10 and FT9/FT10 |
| **Sample rate, Hz** | 207 |
| **Filtering during recording, Hz** | 0.5–48 |
| **Sleep scoring automatic algorithm** | An adjusted to two sqEEG channels version of U-Sleep (Helge et al., 2025) |
| **Questionnaires** | Sleep Diary (Ahrens et al., 2024)<br>Major Depression Inventory (Olsen et al., 2003): mean = $6.2 \pm 2.5$, range = 0 – 20, CV = 0.4<br>World Health Organization 5-item Well-Being Index (Topp et al., 2015): mean = $68 \pm 8$, range = 24 – 100, CV = 0.1 |

*sqEEG – subcutaneous EEG, CV – coefficients of variation*

### EEG

#### Spectral power

To ensure an adequate macrostructural analysis, only nights with at least 5 hours of sleep were included in the analysis. Nights with more than 25% of wake or artifact epochs as defined by the adjusted U-Sleep algorithm (Helge et al., 2025) were excluded. This resulted in 6,429 nights of 20 participants in total (322 nights per participant).

Offline EEG data analyses were carried out with MATLAB (version R2021b, The MathWorks, Inc, Natick, MA), using Fieldtrip toolbox and custom-made scripts. For each participant, we averaged the EEG signal over both electrodes (Table 1), calculated its spectral power for every 30 seconds and differentiated the total power into its aperiodic and oscillatory components using the Irregularly Resampled Auto-Spectral Analysis (Wen & Liu, 2016) as described elsewhere (Rosenblum et al., 2025). Briefly, we transformed the aperiodic power component to log-log coordinates and calculated its slope to estimate the power-law exponent (the rate of spectral decay) in the 1–28Hz range. The oscillatory component was derived by subtracting the aperiodic component from the total power.

#### Metrics

First, as traditional *microscale metrics*, we calculated the relative power of sigma (11 – 16Hz) and slow-wave activity (SWA, 1 – 4Hz). For this, we divided the oscillatory power in the frequency band of interest by the oscillatory power in the broad band (1 – 28Hz). Then, we averaged sigma power over all N2 epochs, and SWA – over either N3 or N2.

The time series of the oscillatory power component in the sigma band (11–16Hz: the frequency range of sleep spindles) was chosen as the hallmark of N2, the sleep stage that comprises up to 50% of sleep. SWA (1–4Hz) was chosen as the hallmark of slow-wave sleep (SWS), the stage that dominates the first half of the night and has several house-keeping physiological functions (Bódizs et al., 2024).

Likewise, we averaged the aperiodic slopes over each sleep stage (Supplemental Fig.1). In addition, we calculated proportions of each sleep stage, REM/NREM ratio and sleep duration derived from sleep stages that were automatically scored with the open-sourced USleep package adapted for sqEEG (Helge et al., 2025).

Second, as a new *macroscale unit* of sleep structure, we derived the time series of smoothened aperiodic slopes (Supplemental Fig.2), SWA or sigma power for each night for each participant (Fig.2). To create these time series, we z-transformed all values of a given night and then smoothened them using the Savitzky-Golay filter as described in our previous work, using the open-source script provided there (Rosenblum et al., 2025).

### Fractal cycles of sleep

As a *mesoscale sleep metric*, we used “fractal cycles” of sleep defined as the time interval during which the time series of the aperiodic slopes descends from the local maximum to the local minimum, with the amplitudes higher than |0.9| z, and then rises back to the next local maximum. A typical night has 4 – 6 fractal cycles that last around 90 minutes and coincide with the NREM-REM sleep cycles defined by hypnogram (Rosenblum et al., 2025). Here, we assessed the following parameters of fractal cycles of sleep: cycle duration, duration of the increasing and decreasing phases of a fractal cycle separately as well as the absolute amplitudes of fractal descent and ascent.

### Dynamic time warping

To quantify how (dis)similar sleep macrostructure was across nights, we calculated the distances between each pair of time series of consecutive nights of either the smoothened aperiodic slopes, SWA or sigma power. For this, we used Dynamic Time Warping (DTW) and, specifically, Matlab’s function *dtw* with its default parameters. DTW temporally aligns two time series by allowing non-linear *warping* (stretching or compression), so that similar patterns can be matched even if they occur at slightly different times (Fig.1, left). After alignment, DTW computes distance between two time series as the sum of Euclidean distances between corresponding points (Salvador & Chan, 2007).

**Figure 1.**
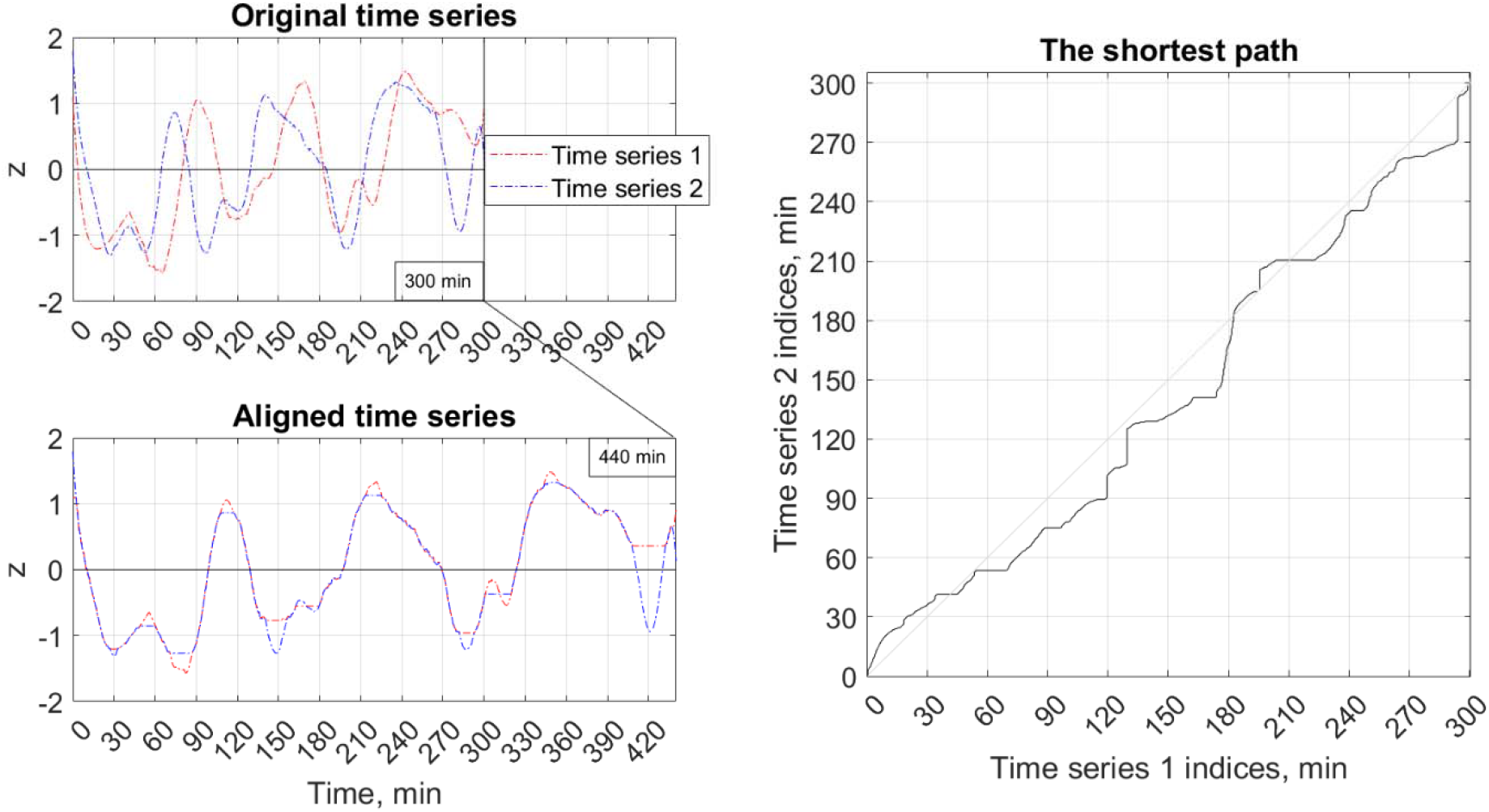
Dynamic Time Warping. Example time series of smoothened, z-scored aperiodic slopes for two consecutive nights: original **(top left)** and aligned **(bottom left)** by the DTW algorithm. For the best alignment, the algorithm warped the time series by 140 min (or 47% of their original length). **Right:** The black line indicates the shortest possible non-Euclidean *path* between the two time series shown on the left. The path shows *how* time series align over time (i.e., the indices where they are warped) and is used to calculate the *distance* between time series (which equalled 43.6 min here), which in turn indicates *how much* they differ after alignment. The distance is further normalized by dividing it by the length of the original time series (300 min) to produce the *dissimilarity metric (which equalled* 14.5% here). The diagonal grey line shows the path between identical time series (no warping, zero distance). A longer path (farther away from the diagonal line) corresponds to higher dissimilarity between two time series. This figure reveals that these two example time series have quite similar macrostructural shape (only 14.5% dissimilarity), which is, however, quite shifted in time (47% of warping needed for alignment). Fig.2 shows *all* time series and dissimilarity histograms in one example participant, Supplemental Fig.2–3 show them for each participant. DTW – Dynamic Time Warping.

Because DTW distances depend on the length of the time series, we truncated all time series to the first 5 hours of sleep (from sleep onset, defined as the first N1 epoch) to ensure equal-length signals (Fig.1, top left). This standardization reduced the number of nights available for macroscale analysis compared to other sleep metrics from 6,429 nights to 5,656 pairs of nights.

The second reason to truncate the analyzed time series at 5 hours stemmed from the fact that the recording device automatically stopped after 6 hours of recording and shortly afterwards automatically started a new recording. Given that one of our aims was to treat the whole-night sleep as an entity, we opted not to use concatenated time series as they had a kink at time of their fusion. The drawback of this decision was that for each night, we excluded an approximately 2-hour episode rich in late-night REM sleep, the period with probable highest inherent variability (e.g., variability related to social factors, such as possible REM rebound on weekends due to late-night REM sleep deprivation on working days when individuals had insufficient sleep duration due to social obligations). At the same time, this decision also had an advantage as the analyzed time series did not include variability related to social obligations and the social jetlag mentioned above and presumably were quite homogeneous.

To ensure comparability, we normalized DTW distances by the length of the time series (5 hours). Normalized distances reflect *the degree of dissimilarity between two time series*, where larger values indicate greater macrostructural differences between nights. Finally, we calculated CV of the dissimilarity metric (as described in the next section) to quantify *intra-* and *inter-individual variation in sleep macrostructure*.

We replicated the DTW analysis using two polysomnographic datasets recorded over three nights and one EEG dataset that recorded up to 15 consecutive nights with a wearable device as described in Supplemental Table 2.

### Coefficients of variation

We assessed stability/variability of sleep metrics with coefficients of variation (CV). CVs are the ratios between standard deviations and means. They are a unitless measure and as such allows comparison between metrics with different units (Winter, 1991).

We calculated CVs of the 1) microscale metrics: sleep stages proportions, sigma power averaged over N2, SWA averaged over N2 and N3 separately, aperiodic slopes averaged over each sleep stage; 2) mesoscale metrics: durations of full fractal cycles of sleep and their descending and ascending parts, and absolute amplitudes of the increasing and decreasing phases of the cycle; 3) macroscale metrics: REM/NREM ratio, sleep duration, DTW distances between time series of sigma, SWA and aperiodic slopes separately (Fig.3).

We compared CVs of different metrics qualitatively by assigning them to one of the four categories: 1) “very low” – CV < 0.1 (i.e., SD is less than 10% of the mean), indicating that a given metric shows high consistency across nights (for *intra-individual* comparison, n ∼ 322 nights/participant) or participants (for *inter-individual* comparison, n = 20); 2) “low” variation across nights/participants – CV = 0.1 – 0.2; 3) “medium” variation across nights/participants – CV = 0.2 – 0.4; 4) “high” – CV > 0.4, indicating low regularity/high unpredictability of a metric. This interpretation is based on previous literature from other areas (e.g., the enzyme-linked immunosorbent assay) (Gill, 1987; Winter, 1991; Reed et al., 2002; Novo et al., 2015; Chouraki et al., 2023) and slightly modified by us to fit it to the EEG area with its relatively low signal-to-noise ratios. The division into four categories is rather arbitrary, yet convenient as in general, lower CV indicates lower standard deviation, less data dispersion and therefore, a certain level of similarity or consistency across nights or equivalence – across participants (Novo et al., 2015) (Fig.3, Supplemental Fig.1).

### Terminology notes

Throughout this article, the word “variation” refers to the *observed* fluctuations in magnitudes of sleep metrics (measured here with CVs), while the word “variability” refers to its *inherent tendency* to fluctuate from night to night, being the inverse of the widely used term “sleep regularity” and is used as a theoretical concept, not as a metric. The word “dissimilarity” refers to *observed* night-to-night variation in macroscale structure (measured here with normalized DTW distances) and is supposed to reflect *inherent* sleep irregularity as a concept.

### Questionnaires

#### Subjective sleep quality

Alongside sqEEG, each morning all participants filled out the sleep diary designed for this study as described in the original paper (see Fig.S2 in Supplemental Material in Ahrens et al., 2024). Here, we used a single item from this diary. Namely, the statement “My sleep was good” rated on a Likert scale from 1 (“Strongly disagree”) to 5 (“Strongly agree”), to measure subjective sleep quality. We averaged this subjective score over each month and correlated it with CVs of all 23 sleep EEG metrics calculated over each month (rather than across the entire year as for the rest of the analyses) pulling all participants together.

#### Well-being and depression

Well-being was measured with the World Health Organization 5-item Well-Being Index (WHO-5; Topp et al., 2015) and Major Depression Inventory (MDI; Olsen et al., 2003) 9 times over a year (approximately once in 1.3 months). The scores were averaged over all measurements across a year and correlated with CVs of EEG metrics calculated over the entire year in all participants together. WHO scores were multiplied by 4 to translate to a percentage scale from 0 (absent) to 100 (maximal) as recommended in the original paper. Higher WHO-5 scores indicate better well-being. Higher MDI scores indicate higher depressive symptomatology. All questionnaires were administered in Danish.

#### Correlations

We used Spearman’s correlation with the Benjamini-Hochberg adjustment to control for multiple comparisons (23 tests) with a false discovery rate set at 0.05 and the α level set in the 0.0022 to 0.05 range.

## Results

### Macroscale unit of sleep structure

We used time series of smoothened z-normalized EEG power (specifically, sigma power, SWA and aperiodic slopes, Fig.2A) as a macroscale unit of sleep structure and assessed night-to-night dissimilarity between these units, using distances calculated by the DTW algorithm. Given that we needed to ensure equal-length signals (Fig.1), for the DTW analysis, we included 5,656 pairs of nights of sleep (283 pairs of nights per participant).

**Figure 2.**
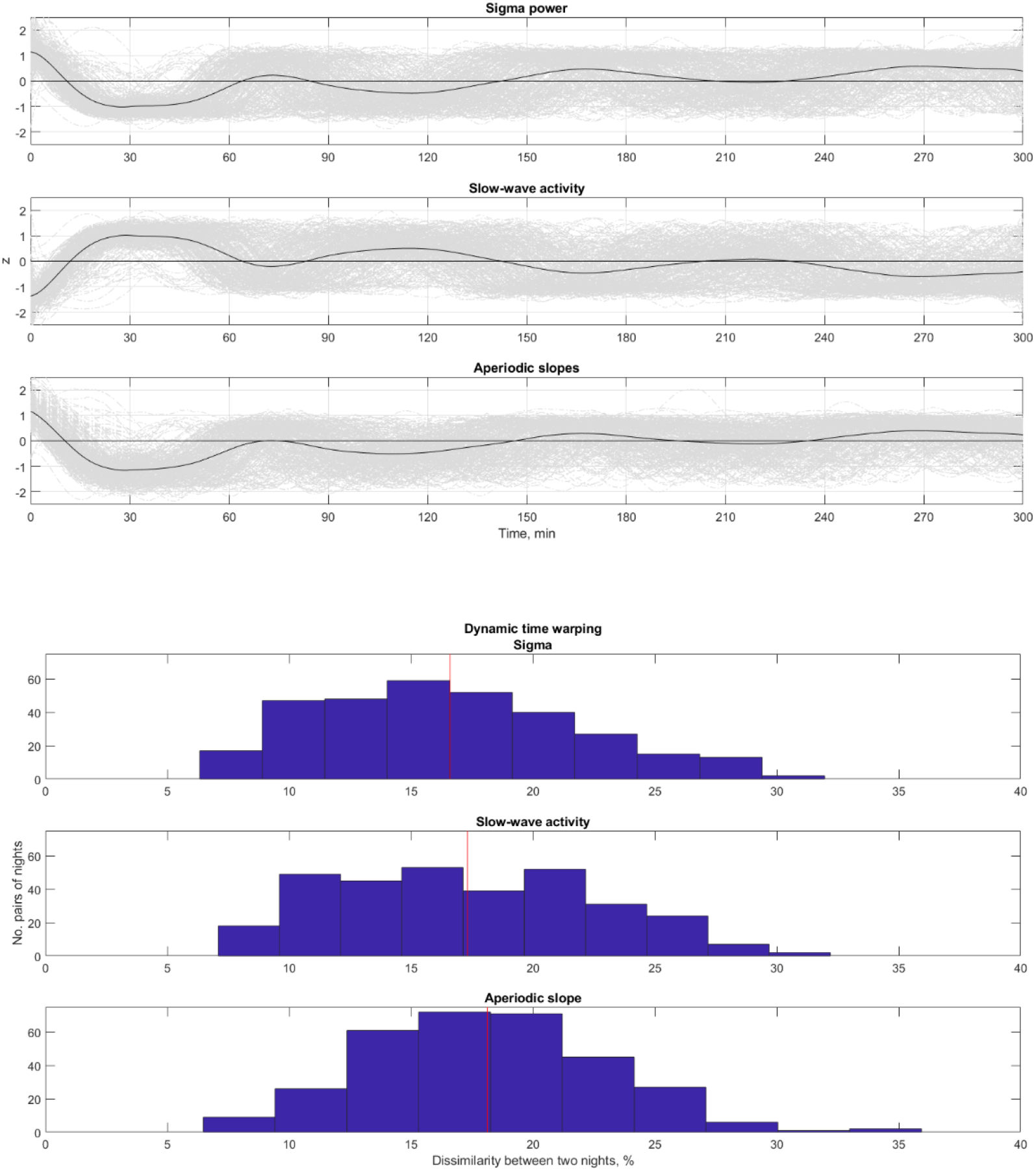
EEG time series as the macroscale unit of sleep structure. **Top**. Time series of z-normalized smoothened sigma power (top), slow-wave activity (middle) and aperiodic slopes (bottom) in one representative participant, starting from sleep onset. The black line shows the time series averaged over all nights of a year (n = 321). The grey lines represent single nights. Supplemental Fig.2 shows such graphs for each participant. **Bottom**. Frequency distribution of dissimilarity between time series of sigma power (top), slow-wave activity (middle) and aperiodic slopes (bottom) shown in A for one example participant. Dissimilarity was calculated by normalizing the DTW distances (mean: 53 – 54 min, SD: 14 – 16 min) by the length of the time series (300 min). The red vertical line shows the mean value, which for this participant, was around 18%, ranging from 6% to 36% for different pairs of adjacent nights. Coefficients of variation in dissimilarity across a year were “medium” (0.26 – 0.32). Supplemental Figure 3 shows histograms for each participant as well as the pooled histogram.

DTW stretches two time series to achieve the best possible alignment between them (Fig.1). On average, 5h-long time series were stretched by 175 ± 16 minutes (i.e., warping amount of 58% of their length) to align them such that peaks and troughs of one nights coincide with the peaks and troughs of the other night, and descending and ascending parts of a cycle – with those of the subsequent night.

After this alignment, the DTW algorithm estimates the distance or path between two time series. The longer the path, the larger the dissimilarity between them. We found that the mean distance between two adjacent 5h-long time series was around 55 – 59 minutes, i.e., 18 – 20% of their length, ranging intra-individually from a minimum of 5% to a maximum of 75% (Fig.2B, Supplemental Fig.3). In other words, on average, the macroscale sleep structure of two adjacent nights showed ∼20% dissimilarity.

To summarize, night-to-night macrostructure dissimilarity looks quite low, only ∼20%, yet we should keep in mind that before this computation, the algorithm first warped the time series by 58%. These two metrics should be taken and interpreted together, indicating that while the overall macrostructural patterns are relatively similar across nights, their temporal alignment is highly variable.

### Variation in sleep metrics

Table 2 reports the means of intra-individual means, SD and CVs as well as inter-individual CVs of all metrics. Fig.2 shows intra-individual CVs of all sleep metrics for each participant individually. We compare CVs of different metrics qualitatively only as belonging to either the “very low”, “low”, “medium” or “high” categories.

**Table 2.**
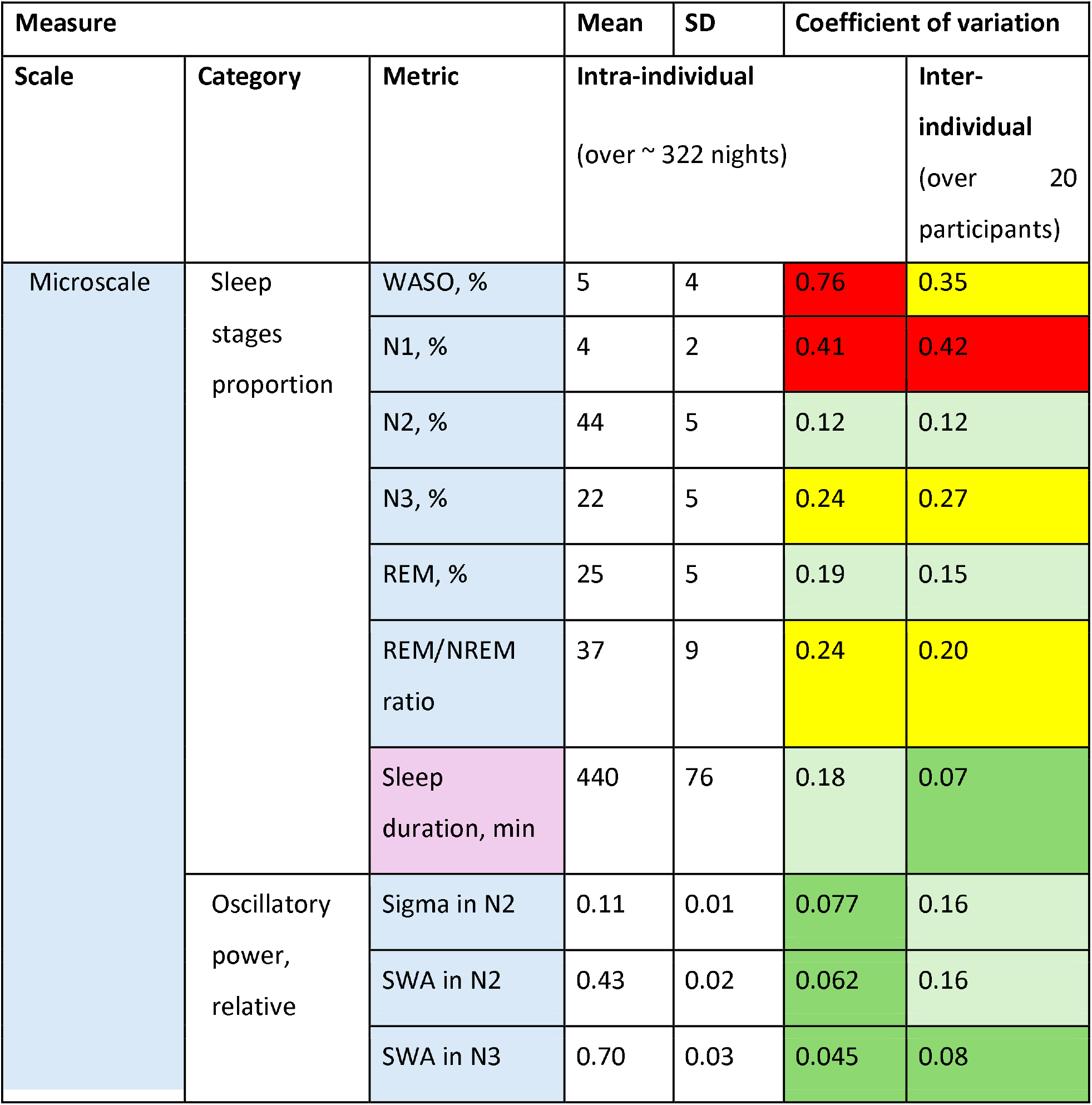
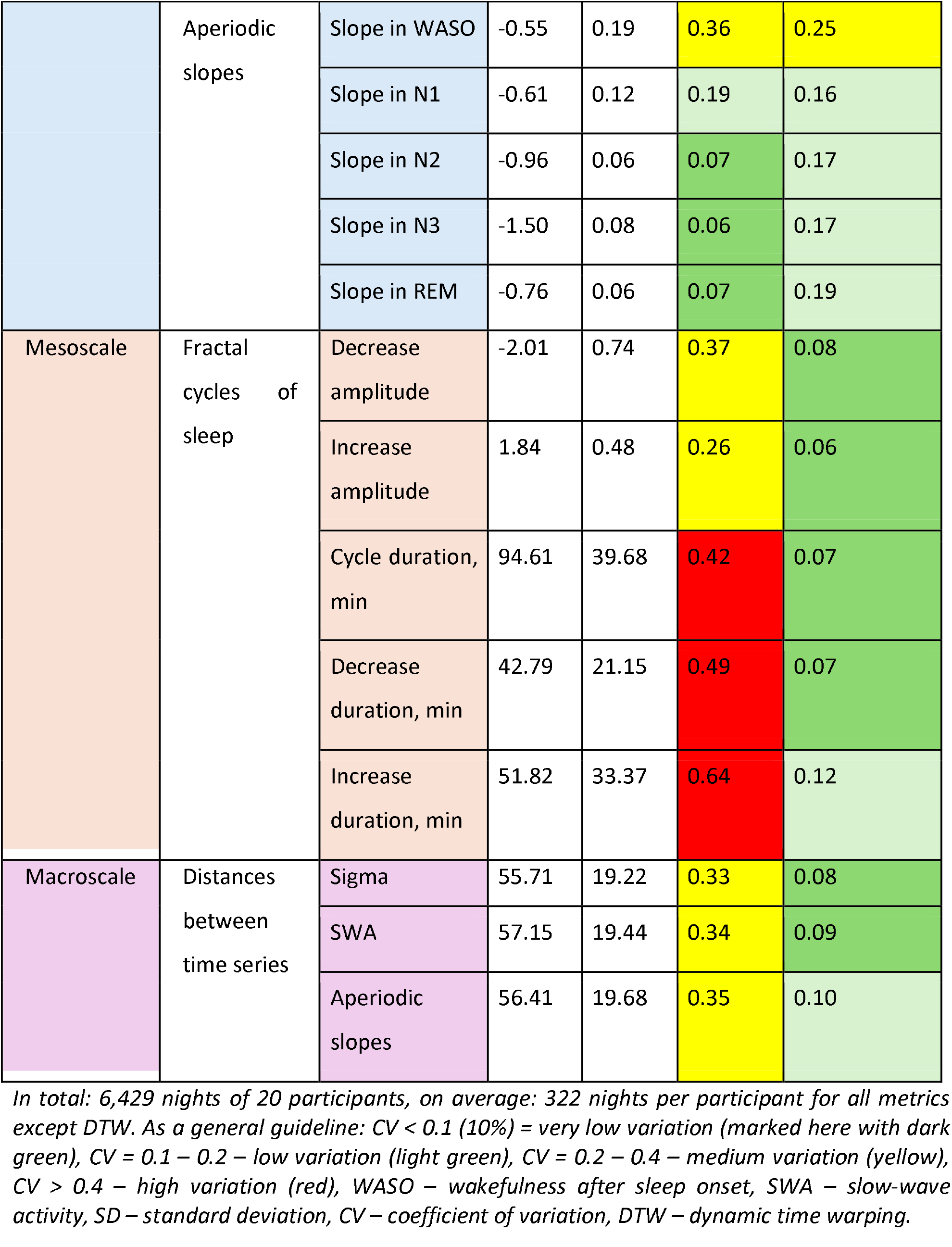
Sleep metrics: mean, SD, CV.

### Microscale metrics – spectral power and sleep stages

Spectral power, namely, its oscillatory component in the sigma and SWA bands during N2 and N3 and the slopes of its aperiodic component during N2, N3 and REM sleep showed “very low” *intra-individual CV* (range: 0.05 – 0.08), indicating that the data is highly consistent from night-to-night. At the *inter-individual* level, most of these metrics showed “low” *CV*, except for SWA in N3, which was “very low” (Table 2). Aperiodic slopes averaged over WASO showed “medium” intra- and inter-individual CV.

Sleep stage proportions, REM-NREM sleep ratio and TST showed “low” to “medium” intra-individual CV (range: 0.12 – 0.41), except for WASO that showed “high” CV (Fig.3). Inter- and intra-individual CV of these metrics were comparable, except for sleep duration that showed “very low” variation (Table 2).

### Mesoscale metrics – fractal cycles of sleep

We quantified sleep cycles, using the recently introduced concept of “fractal cycles of sleep” based on temporal fluctuations of aperiodic slopes across a night (Rosenblum et al., 2025). Given that for consistency between nights and subjects, we assessed only the first 5h of sleep, there were only 2 – 3 sleep cycles per night per participant. Fractal cycle durations showed “high” *intra-individual* CV (range: 0.42 – 0.64), where CV of the duration of the increasing part of the cycle was higher than that of the decreasing part of the cycle and the whole cycle. *Intra-individual* CV of the amplitudes of the fractal decrease and increase were “medium”. At the same time, *inter-individual* CV of all sleep cycle parameters were “very low”, except for duration of cycle increase that was “low” (Fig.3, Table 2).

### Macroscale metrics – time series dissimilarity

Distances between successive time series as a measure of dissimilarity of macroscale sleep structure showed “medium” *intra-individual* and “very low” *inter-individual* CVs (Table 2, Fig.3).

**Figure 3.**
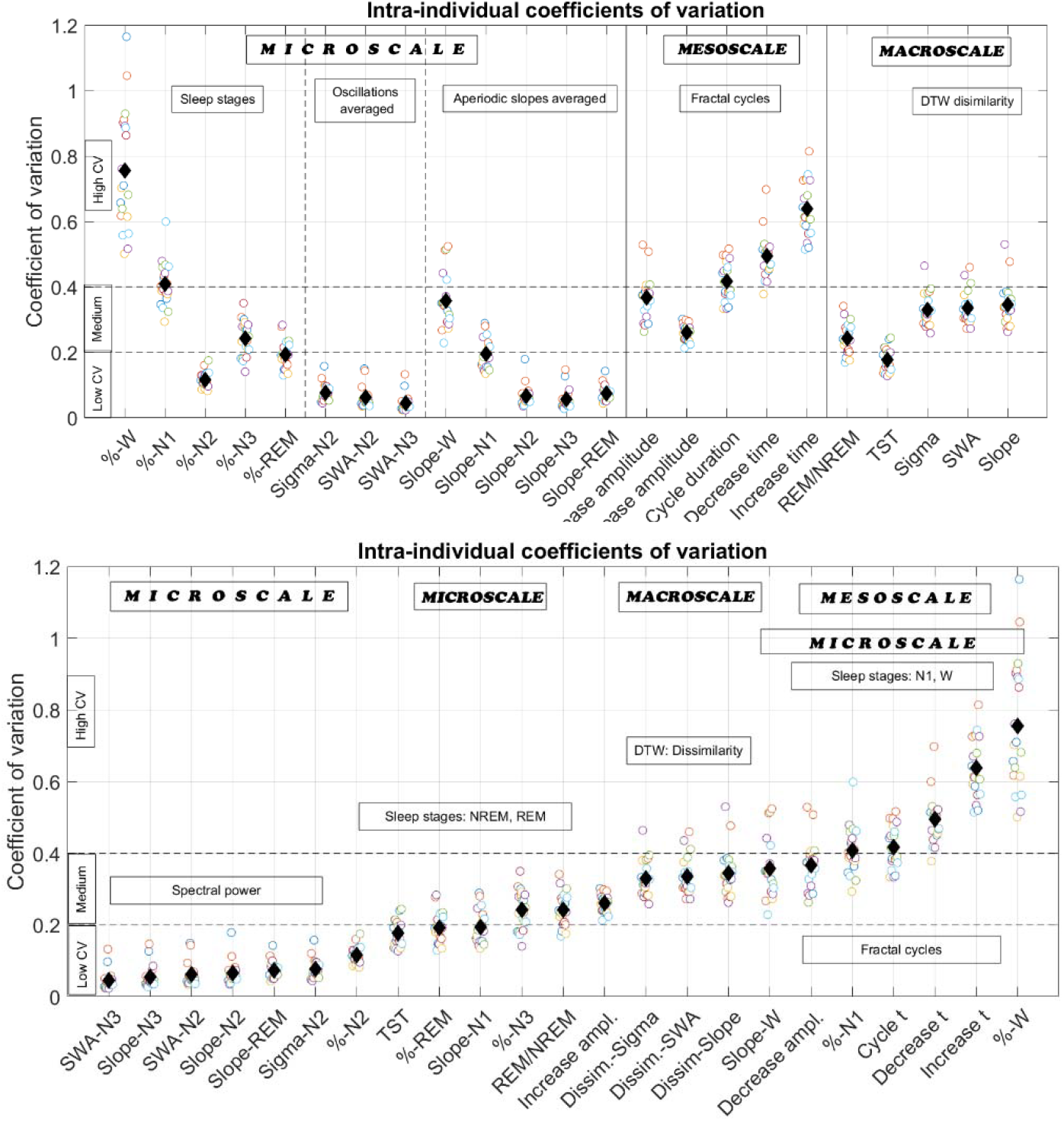
Intra-individual coefficients of variation of sleep metrics. Each open circle represents the CV (y-axis) of a given sleep metric (x-axis) in one individual; n = 20, thus, 20 circles for each metric. Black diamonds show CV averaged over all participants. CVs of the following traditional and new sleep metrics are shown: 1) microscale level: sleep stages proportions, sigma power averaged over N2, SWA averaged over N2 and N3, aperiodic slopes averaged over each sleep stage; 2) mesoscale level: amplitudes (“ampl.”) of the decrease and increase of fractal cycles, durations (“t”) of the whole fractal cycles and their descending and ascending parts; 3) macroscale level: REM/NREM ratio, TST, DTW dissimilarity (see Fig.2) between time series of sigma power, SWA, and aperiodic slopes. On average, 322 nights per participant were used for each metric except DTW (where 283 pairs of nights > 5h long were available on average). CV is the ratio of the standard deviation to the mean of all values of a given year. As a general guideline: CV < 0.1 – very low variation, CV = 0.1 – 0.2 – low variation, CV = 0.2 – 0.4 – medium variation, CV > 0.4 – high variation. **Top**. Sleep metrics sorted by their scale: from micro-to meso- and macroscale. **Bottom**. Sleep metrics sorted by their CV: from low to medium and high. WASO – wakefulness after sleep onset, TST – total sleep time, REM – rapid eye movement sleep, SWA – slow-wave activity, DTW – dynamic time warping, CV – coefficient of variation, t – time, ampl. – amplitude.

### Correlations with subjective sleep quality

We correlated between CV of all EEG sleep metrics and subjective sleep quality within all participants. Lower variation in N1 proportion and aperiodic slope amplitudes during N1 and WASO, as well as in sleep structure dissimilarity (as assessed by distances between time series of either sigma, SWA or aperiodic slopes) correlated with higher subjective sleep quality. Higher variation in N3 proportion as well as sigma oscillations, SWA and aperiodic slopes during N2 and N3 correlated with higher subjective sleep quality (Table 3, Fig.4). All significant correlations were weak (r < 0.3). Of note, except for N3, metrics with “very low” CVs showed positive correlations, while metrics with “medium” CVs showed negative correlations.

**Table 3.** Correlations between CV of objective sleep metrics and subjective sleep quality.

| Scale | Category | Metric | r | p |
| --- | --- | --- | --- | --- |
| Microscale | Sleep stages proportion | WASO | 0.070 | 0.244 |
|  |  | N1 | -0.209 | 0.0004 |
|  |  | N2 | -0.058 | 0.334 |
|  |  | N3 | 0.229 | 3E-4 |
|  |  | REM | -0.002 | 0.977 |
|  |  | REM/NREM ratio | 0.051 | 0.394 |
|  |  | Sleep duration | 0.102 | 0.089 |
|  | Oscillatory power | Sigma in N2 | 0.198 | 0.001 |
|  |  | SWA in N2 | 0.297 | 3E-7 |
|  |  | SWA in N3 | 0.262 | 3E-4 |
|  | Aperiodic slopes | Slope in WASO | -0.140 | 0.018 |
|  |  | Slope in N1 | -0.247 | 0.000 |
|  |  | Slope in N2 | 0.290 | 3E-5 |
|  |  | Slope in N3 | 0.247 | 3E-5 |
|  |  | Slope in REM | -0.052 | 0.387 |
| Mesoscale | Fractal cycles of sleep | Decrease amplitude | 0.114 | 0.057 |
|  |  | Increase amplitude | -0.120 | 0.044 |
|  |  | Cycle duration | -0.039 | 0.513 |
|  |  | Decrease duration | 0.107 | 0.072 |
|  |  | Increase duration | -0.036 | 0.543 |
| Macroscale | Macrostructure<br>dissimilarity | Sigma DTW | -0.249 | 7E-05 |
|  |  | SWA DTW | -0.252 | 6E-05 |
|  |  | Aperiodic slopes<br>DTW | -0.153 | 0.016 |

*Spearman's correlations between CVs of the objective sleep EEG metrics calculated over each month and subjective sleep quality based on the Likert scale assessment of the statement "My sleep was good", ranging from 1 ("Strongly disagree") to 5 ("Strongly agree") recorded each morning and averaged over each month. P-values adjusted with the Benjamini-Hochberg correction for 23 comparisons with the $\alpha$ level set in the 0.0022 to 0.05 range. $r$ – Spearman's correlation coefficient, $r < 0.3$ are considered as weak, $r = 0.3 - 0.7$ – medium, and $r > 0.7$ strong correlations. WASO – wakefulness after sleep onset, TST – total sleep time, REM – rapid eye movement sleep, SWA – slow-wave activity, DTW – dynamic time warping, CV – coefficient of variation.*

**Figure 4.**
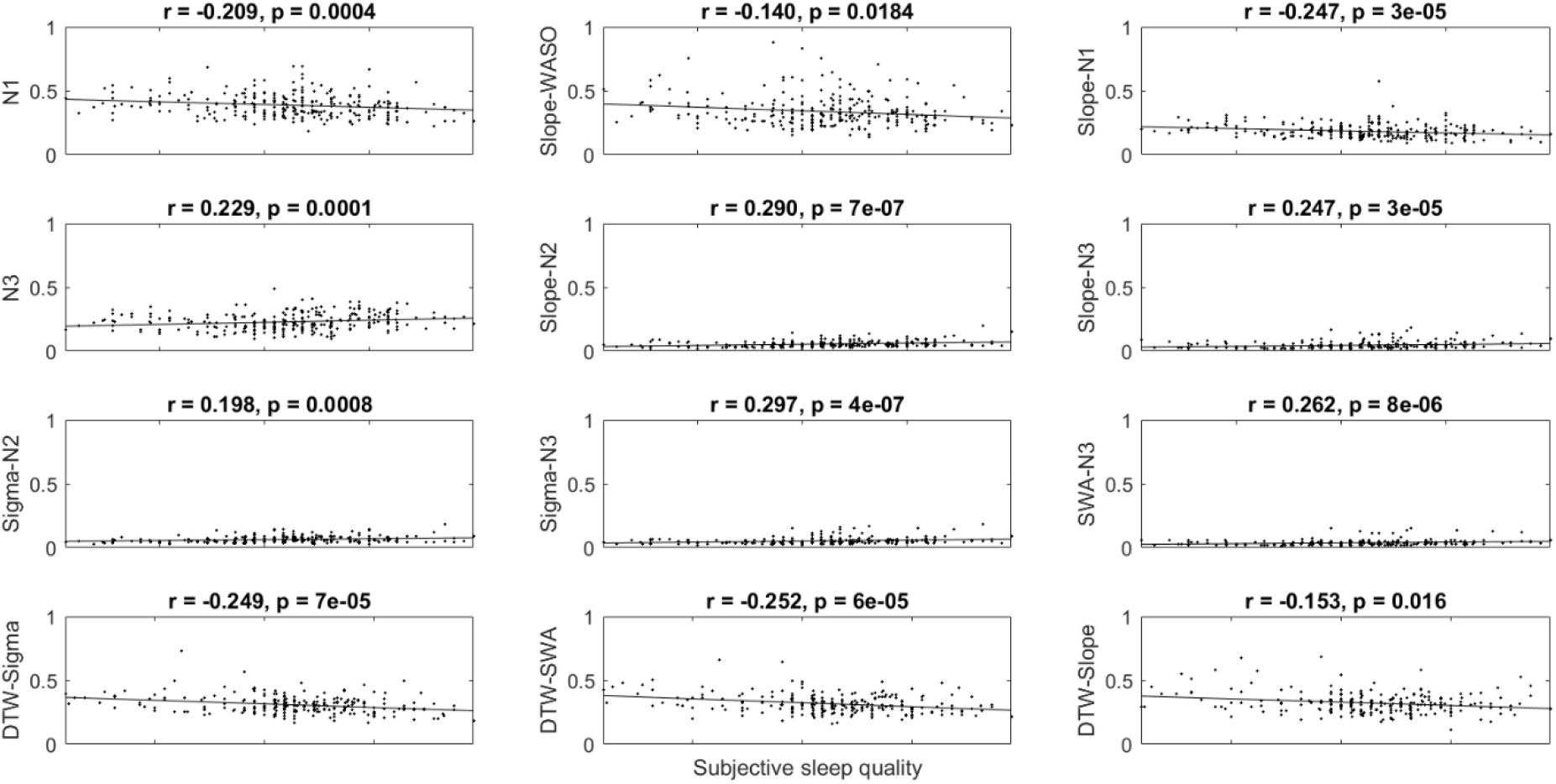
Correlations between CVs of objective sleep metrics and subjective sleep quality. Only the significant Spearman’s correlations reported in Table 3 are shown here. Sleep metrics were calculated over each month of the year and pulled for all participants. **First row:** Lower variation in N1 proportion and aperiodic slopes in WASO and N1 correlated with higher subjective sleep quality. **Second row:** Higher variation in N3 proportion and aperiodic slopes during N2 and N3 correlated with higher subjective sleep quality. **Third row:** Higher variation in sigma in N2 and SWA during N2 and N3 correlated with higher subjective sleep quality. **Fourth row:** Lower variation in sleep structure dissimilarity (as assessed by DTW distances between time series of either sigma, SWA or aperiodic slopes) correlated with higher subjective sleep quality. All correlations were weak (all r’s < 0.3). Except for N3, objective metrics with “very low” CVs (Table 2) showed positive correlations, while metrics with “medium” CV showed negative correlations. r – Spearman’s correlation coefficients, WASO – wakefulness after sleep onset, SWA – slow-wave activity, DTW – dynamic time warping, CV – coefficient of variation.

### Correlations with subjective well-being

Subjective well-being and CVs of EEG sleep metrics calculated over one year did not correlate after the correction for multiple comparisons (all p’s > 0.002), indicating that there is either no association or that these questionnaires are not sensitive enough (a possible “floor effect”) to detect possibly existing associations in healthy participants with overall very good estimation of self-well-being.

## Discussion

Using the longest continuous healthy sqEEG dataset to date (Ahrens et al., 2024), we introduced the macroscale metric of whole-night sleep and measured its night-to-night dissimilarity. Based on the analysis of these new metrics, we conclude that while the overall macrostructural patterns of sleep are relatively similar across nights (as reflected by dissimilarity of ∼20%), their temporal alignment is quite variable (as reflected by warping amount of ∼60%). We further compared yearlong variation in macrostructure with that of the conventional micro- and mesoscale metrics and found that different sleep parameters showed more metric-specific than scale-dependent variation.

### Macroscale: Whole-night sleep as an entity

Visually, EEG time series might look quite different due to differences in the timing of peaks and troughs, and durations of descending and ascending parts of time series, yet after alignment, they often show noticeable similarity (Fig.1). The DTW analysis captured this dissociation directly: the residual dissimilarity after alignment was small (20%), whereas the warping required to align nights was relatively large (58%). This observation suggests that night-to-night differences in sleep macrostructure are predominantly temporal rather than compositional.

Sleep macrostructure showed medium night-to-night *intra-individual* variation (CV∼0.35), indicating that each night contains both stable, trait-like patterns repeated from night to night as well as random and unpredictable ones. Interestingly, *inter-individual* variation was “low” (CV∼0.1), meaning that the data was consistent over the participants and they all presented with “medium” *intra-individual* night-to-night variation (Fig.2). Importantly, we were able to replicate our findings in two independently collected datasets (Supplemental Table 2).

A recent study by Leguia et al. (2025) who also used sqEEG and DTW in healthy participants, but over 30 days, found lower within-than between-subject dissimilarity. Of note, the aim of this study was to compare CVs of different metrics, and not their means as in Leguia et al. Nevertheless, for the replication purpose, we performed a supplemental analysis to compare within- and between-subject dissimilarity and replicated the previous finding.

### Sleep regularity

In view of these results, we suggest that the measure of macrostructure dissimilarity introduced here can be utilized as a novel sleep regularity metric. (We have to note here that even though linguistically, the commonly used in sleep research word “regularity” is the opposite of “dissimilarity”, mathematically, 20% “dissimilarity” measured here does not imply that there was 80% “similarity”, since this has not been quantified and compared directly thus far. We hope that future mathematical studies will quantify the link between these two concepts directly.)

Several metrics of sleep regularity have been used in research thus far, including inter-daily stability index, a social jetlag measure, composite phase deviation, the sleep regularity index as well as intra-individual standard deviation and coefficient of variation of such metrics as sleep onset/offset time, mid-sleep time and sleep duration (reviewed by Hartstein et al., 2025). These metrics are mostly derived from actigraphy or self-reports, which lack direct access to brain activity, and therefore, capture regularity of sleep timing/schedule only. In this context, the macrostructure variation metric introduced here may serve as a complementary measure of sleep regularity, quantifying the stability/variability of whole-night neural sleep organization across nights.

Sleep regularity is an important characteristic of sleep. Poor sleep regularity has been linked to perceived stress and depressed mood (Lunsford-Avery et al., 2018; Pye et al., 2021; Magal et al., 2022). Strikingly, low sleep regularity showed stronger mortality risk than sleep duration (Windred et al., 2024). In line with the direction of this literature, we observed that lower variation in sleep macrostructure correlated with better subjective sleep (Fig.4). On the other hand, it did not correlate with subjective well-being, yet this might reflect the “ceiling effect” as all participants reported good well-being and mental stability (Table 1).

### Mesoscale variation: fractal cycles of sleep

We found that *intra-individual* night-to-night variation in duration of fractal cycles of sleep (used as a proxy of NREM-REM cycles) was “high”. This is in line with the fact that much warping (58%) was needed to align two time series of aperiodic slopes (which comprise fractal cycles). High variation presumably reflects the well-established heterogeneity of individual sleep cycles in terms of durations of their NREM vs REM sleep phases and their substages (i.e., N1, N2, N3 for NREM sleep and tonic-phasic dichotomy for REM sleep). At the same time, the amplitudes of the fractal cycle descents and ascents showed “medium” variation across nights.

Similarly to the macroscale metric, here too, *inter-individual* variation was “low” (CV∼0.1), meaning that “medium” to “high” night-to-night variation was consistently observed in all participants, i.e., they all presented with heterogeneous sleep cycles (Fig.3).

To summarize, the structure of different sleep cycles is quite variable both across one night (Le Bon, 2020) and across different nights of the same individual.

### Microscale variation: spectral EEG power

Sigma oscillations averaged over N2, SWA averaged over N2 and N3, and aperiodic activity averaged over N2, N3 and REM sleep showed “very low” *intra-individual* night-to-night variation. This might reflect the procedure of averaging over hundreds of epochs of a single night, which as such eliminates the temporal dynamics of spectral power and smoothens its richness, providing a stable metric. Our supplemental analysis partly supports this interpretation, showing that variation of spectral power metrics increases almost twice, when calculated over all epochs of a given sleep stage of a given night without averaging. Yet it still remains in the category “low variation” (<20%). This is in line with previous studies reporting that NREM sleep EEG measures, especially, sigma power, show high overnight and across night stability and are probably a genetically determined trait, with a heritability estimate of up to 96% for spindles, which is not influenced by sleep need and intensity (Buckelmüller et al. 2006; de Gennaro et al. 2008; Purcell et al., 2017). Taken together, these findings suggest that sleep microstructure as reflected by spectral power during N2, REM sleep and especially N3, is quite stable not *only* due to averaging. Higher stability of N3 compared to N2 and REM sleep could be further explained by the widely reported observation that the N3 signal is characterized by lowest complexity and entropy due to long-range integration and global synchronization processes (Miscovic et al., 2018). Highly ordered, regular signals with low information content typically seen during N3 probably leave little dynamic range for variability.

Interestingly, *inter-individual* variation was “low” yet still twice as high as *intra-individual* CV, except for SWA in N3, where it was “very low”. Our findings are in line with the sqEEG study by Leguia et al. (2025) who showed that “core dynamics in sleep oscillations are consistently shared within individuals, yet are rather variable over consecutive nights between the subjects”.

Another interesting observation is that metrics with “very low” CVs correlated positively with subjective sleep quality, while metrics with “medium” CVs correlated with it negatively (Table 4). This might mean that small variations in sleep microstructure have some physiological relevance, yet this must be directly tested by future research.

### Variation in sleep stages

N2 and REM sleep and total sleep duration showed “low” *intra-individual* night-to-night variation, whereas variation in N3 and REM/NREM ratio was “medium”. Variation in N1 and WASO was “high” in line with the fact that healthy people spend little time in WASO and N1. However, their proportions might increase substantially on nights with internal (e.g., pain) or external (e.g., noise) disturbances, thereby increasing the SD and CV. *Inter-individual* CVs varied from “low” to “high” (Table 2). These findings are in line with Chouraki et al. (2023) who also showed a metric-specific manner of night-to-night variation.

## Conclusions

Technical and methodological limitations of the Ultra-Long-Term Sleep project are described in its original papers (Djurhuus et al., 2023; Ahrens et al., 2024). This study used that project’s data to introduce new measures of sleep macrostructure and its variability and showed that while the overall macrostructural patterns are relatively similar across nights, their temporal alignment is quite variable. We further showed that different sleep metrics presented a differential degree of night-to-night stability, which was more metric-specific than scale-dependent. This might reflect the distinction between more static, trait-like versus more dynamically varying features of sleep.

We expect that in-detail description of multilevel sleep structure and its stability/variability would allow the extraction of electrophysiological fingerprints, which in turn would allow setting up personalized time-sensitive recommendations to help people facing (social) jet lag or shift work. One specific example application would be personalized timing of alarms that considers individual sleep structure dynamics using metrics with intrinsic very low variability (such as, for example, spectral power here). Multilevel description of sleep structure with disentangling between its trait-like (e.g., heritable) vs state-like patterns might also have clinical implication, where identifying deviations from normal patterns would advance diagnosis and monitoring of sleep disorders, prediction of nocturnal seizures in epilepsy management care, instability in breathing control in apnea etc.

## Acknowledgments

We would like to thank the team who conducted the Ultra-Long-Term Sleep study and the study participants for their time, effort and dedication.

## Funding

MD, LB and YR are supported by the Dutch Research Council (NWO). YR is also supported by the Alzheimer Nederland Early Career Grant. The funders had no role in the design, data collection, data analysis, and reporting of this study.

## Disclosure Statement

Nothing to disclose.

## Data availability

The current study generated no original data, the data source is described in the study design paper (Ahrens et al., 2024).

## Notes

### Competing Interest Statement

The authors have declared no competing interest.

