## Supplemental Material for "Night-to-night sleep EEG variability over one year"

**Part 1: Supplemental to the main dataset**

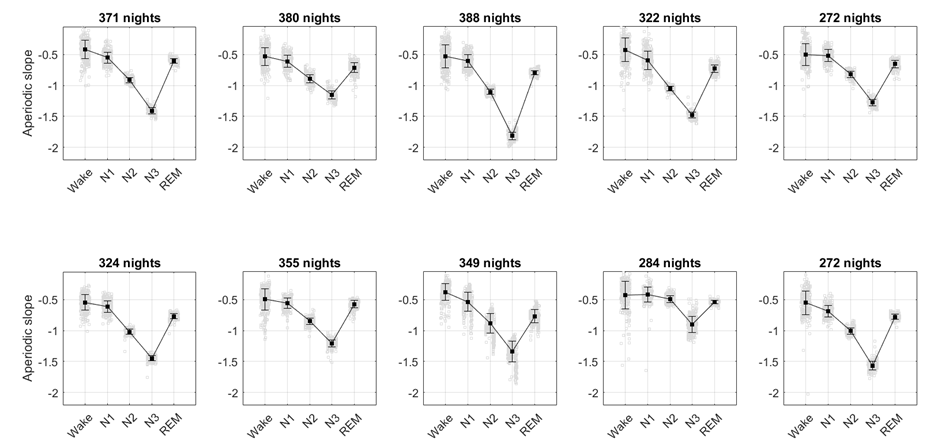

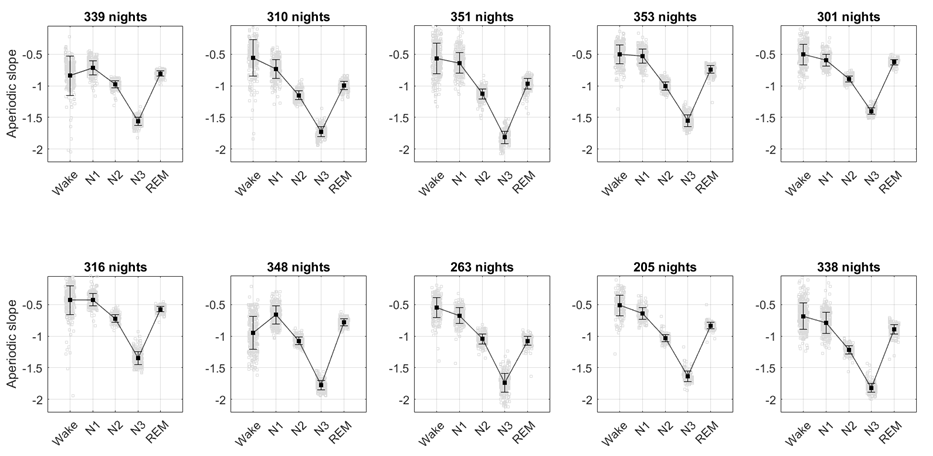

**Supplemental Figure 1. Night-to-night variation in aperiodic slopes.** Each graph shows aperiodic slopes (1–28 Hz) in one participant (n = 20): the grey squares show individual night values, and the black squares show the values averaged across all nights of a year. Aperiodic slopes during N2, N3 and REM sleep show very low intra-individual night-to-night variation as reflected by the small dispersion of grey squares. The coefficients of variation are shown in Fig.3 and Table 2.

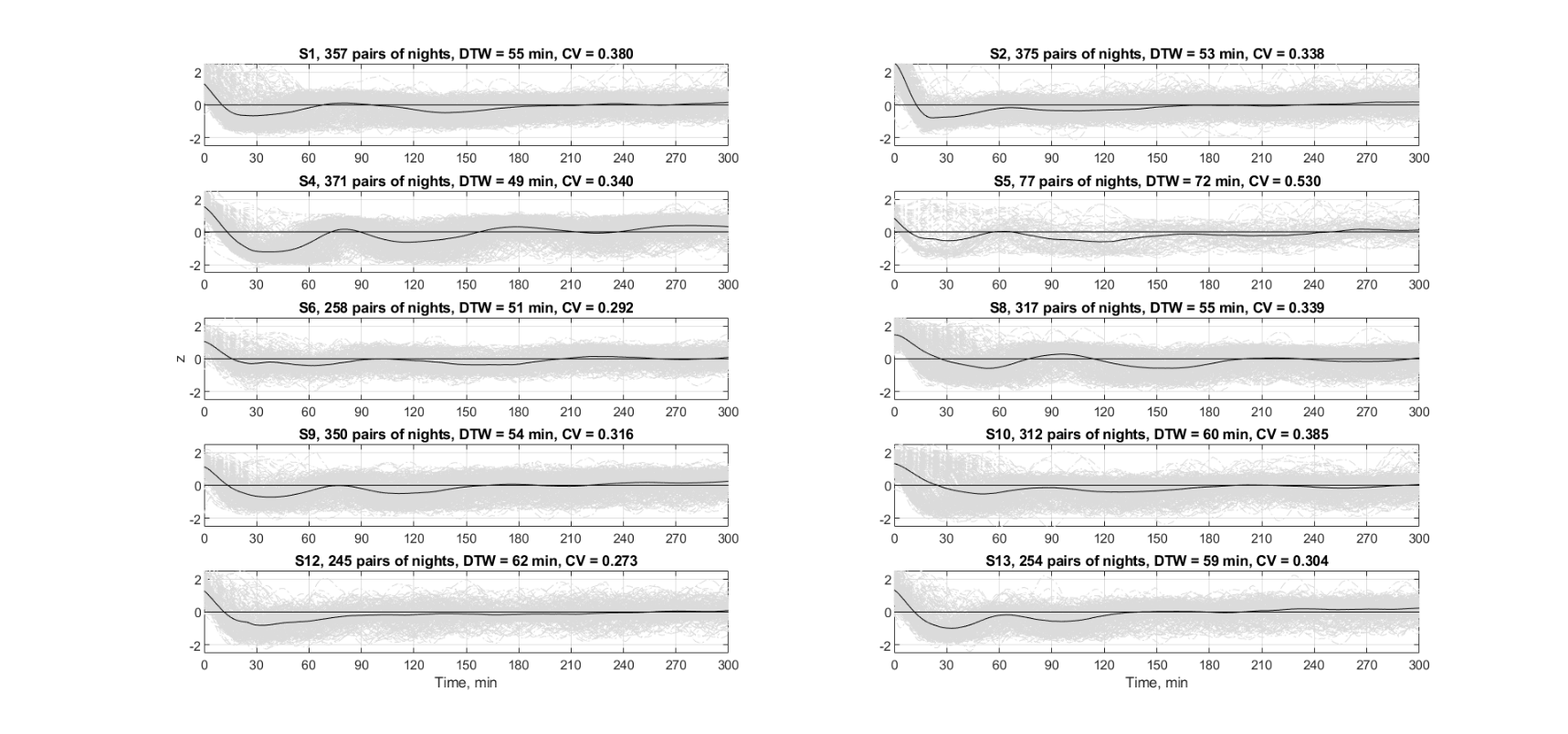

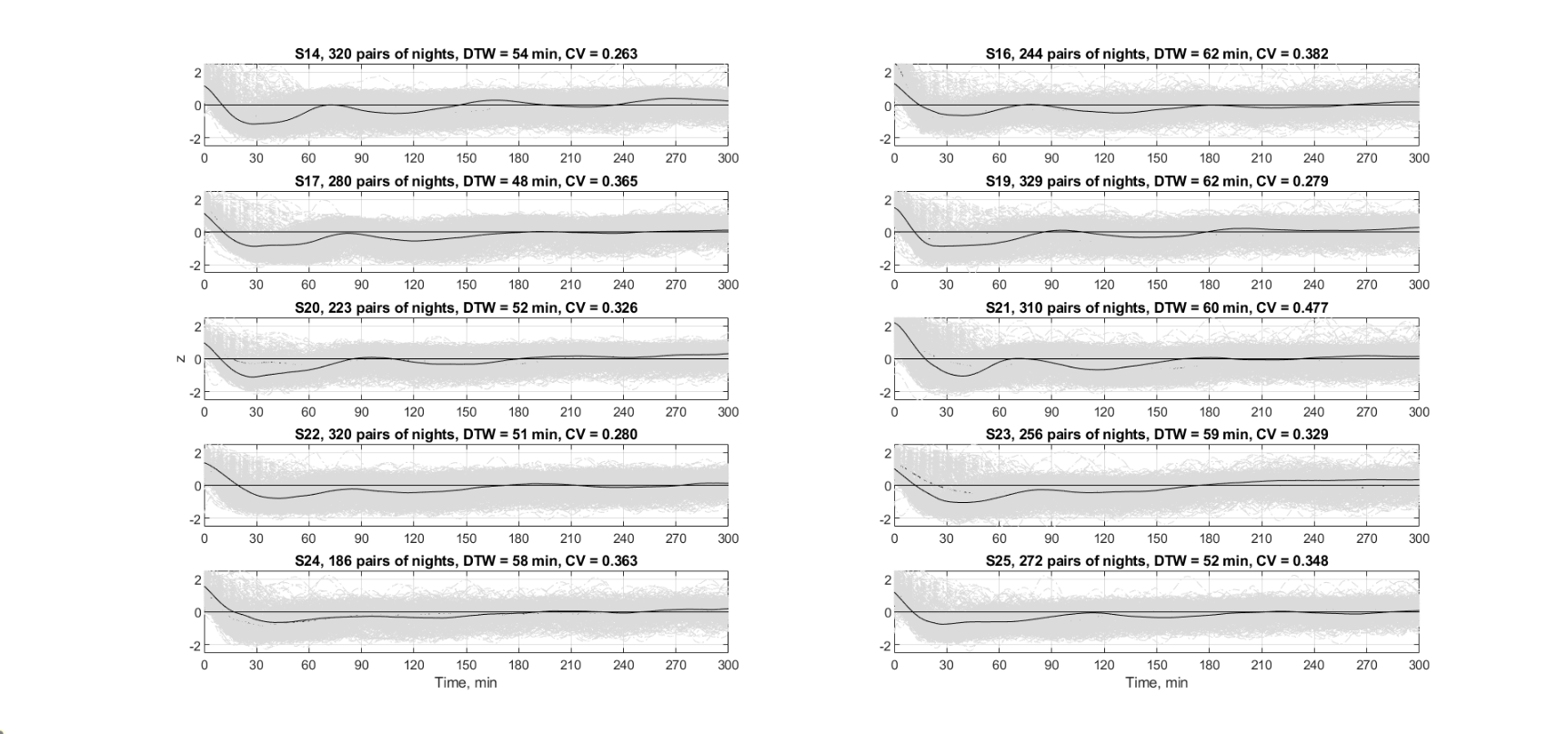

**Supplemental Figure 2. The macroscale unit of sleep: Main sqEEG dataset.** Each graph shows a 5h-long time series of z-normalized smoothened aperiodic slopes in one participant (n = 20). Grey lines show time series of individual nights across one year, while black lines reflect the time series averaged over all nights of a given participant. DTW is short for the mean DTW distances between two time series, where a larger distance corresponds to a higher dissimilarity between them. All CVs – the ratios between the standard deviation to the mean distances – were “medium”. DTW – dynamic time warping, CV – coefficient of variation.

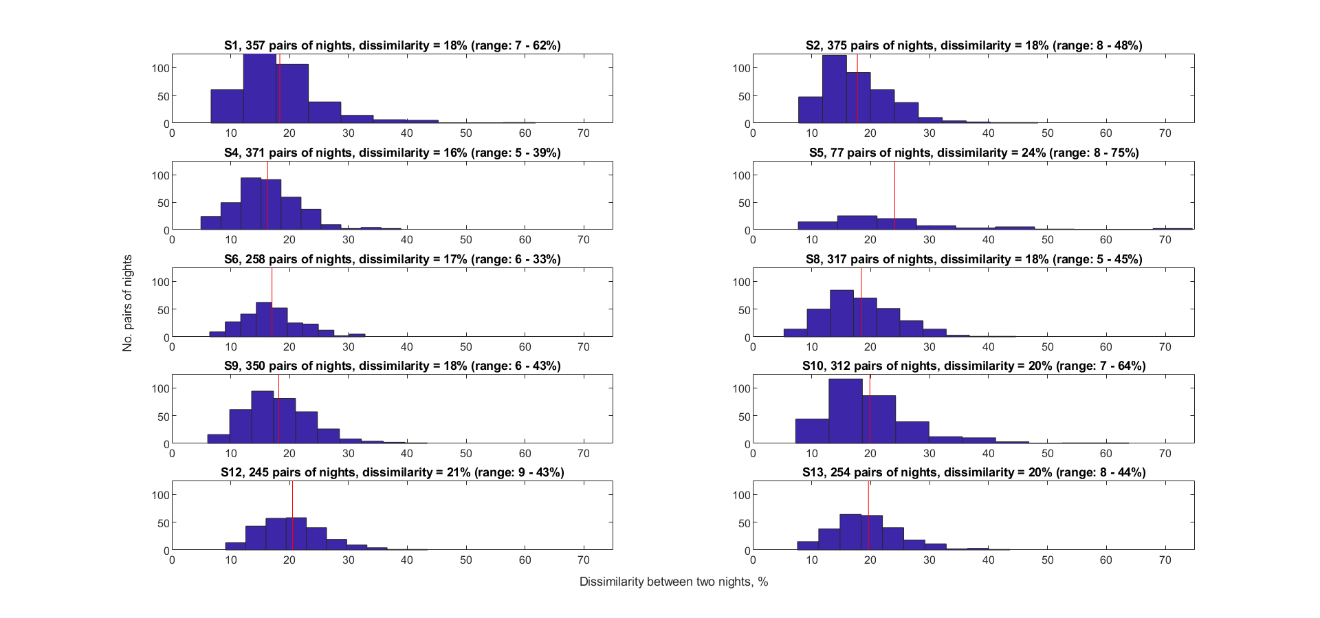

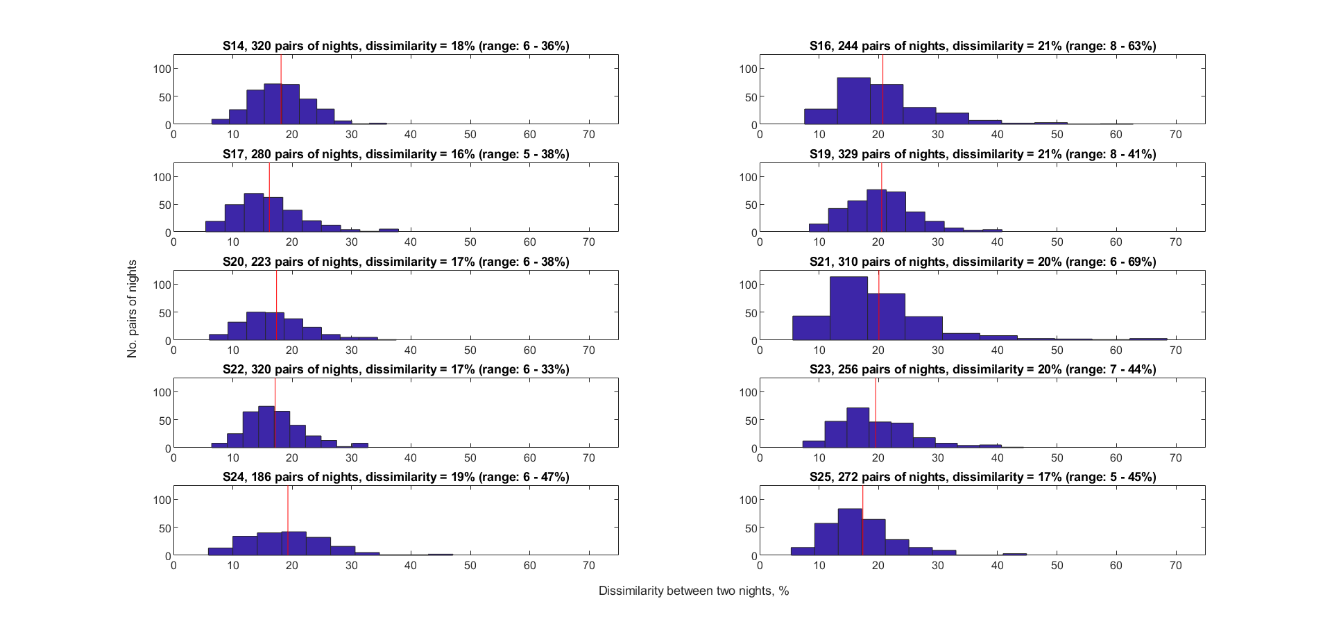

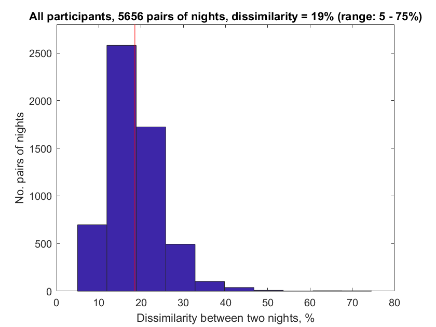

**Supplemental Figure 3. Histograms of intra-individual dissimilarity between time series of aperiodic slopes.** Dissimilarity is the DTW distance (S.Fig.2) normalized by the length of the time series (300 min). Red vertical lines show the mean values. **Top:** Each graph shows a dissimilarity histogram in one participant (n = 20, on average, 283 pairs of nights per participant). **Bottom:** Dissimilarities in all participants pooled together (5,656 pairs of nights). DTW – dynamic time warping.

**Supplemental Table 1: Coefficients of variation of spectral power metrics**

| **Scale** | | **Microscale (individual events)** | | | **Macroscale (whole-night sleep as a unit, DTW)** | |
| --- | --- | --- | --- | --- | --- | --- |
| **Coefficients of variation over** | | **All epochs of a night** | **All nights of a participant** | **All participants** | **All nights of a participant** | **All participants** |
| n | | 43 – 413 | 322 | 20 | 283 | 20 |
| Sigma power in N2 | 413 | 0.113 | 0.077 | 0.133 | 0.330 | 0.082 |
| SWA in N2 | 413 | 0.104 | 0.062 | 0.137 | 0.336 | 0.089 |
| SWA in N3 | 206 | 0.161 | 0.045 | 0.072 |  |  |
| Aperiodic slope in WASO | 48 | 0.308 | 0.358 | 0.252 | 0.345 | 0.100 |
| Aperiodic slope in N1 | 43 | 0.307 | 0.194 | 0.156 |  |  |
| Aperiodic slope in N2 | 413 | 0.130 | 0.066 | 0.170 |  |  |
| Aperiodic slope in N3 | 206 | 0.217 | 0.056 | 0.167 |  |  |
| Aperiodic slope in REM | 222 | 0.180 | 0.074 | 0.191 |  |  |

*In total, 6,429 nights of 20 participants, on average, 322 nights per participant for all metrics except DTW. For DTW, in total, 5,656 pairs of nights, on average, 283 pairs of nights per participant; n refers to either the number of epochs of a night, the number of nights or the number of participants. As a general guideline: CV < 0.1 (10%) = very low variation (dark green), CV = 0.1 – 0.2 – low variation (light green), CV = 0.2 – 0.4 – medium variation (yellow), CV > 0.4 – high variation. WASO – wakefulness after sleep onset, SWA – slow-wave activity, REM – rapid eye movement sleep, N – non-REM sleep, CV – coefficient of variation, DTW – dynamic time warping.*

**Part 2: Replication of DTW in independent datasets**

We replicated the DTW analysis, using the following independently collected datasets (Supplemental Table 2):

Replication Dataset 1: Donders 2022 dataset from a home-based sleep study exploring simultaneous polysomnography and EEG wearables conducted at the Donders Institute for Brain, Cognition and Behavior, the Netherlands (Described as Dataset 2 in Jafarzadeh Esfahani et al., 2023). The signal was recorded at participants’ homes over three nights with a gap of a week between each recording. We used the average of F3 and F4 (Fig.S4). The EEG processing is described in Table 6 in Rosenblum et al. (2025) as Dataset 4. In addition, this dataset includes the signal recorded at participants’ homes over up to 15 consecutive nights with a wearable device (Zmax, Hypnodyne, Sofia, Bulgaria) with two latero-frontal channels and a fronto-central reference (Jafarzadeh Esfahani et al., 2023) (Fig.S5).

Replication Dataset 2: the continuous multi-night sleep database for exploring sleep structure similarity recorded over three consecutive nights conducted at the University of Electronic Science and Technology of China. We included only participants who had three recordings with WASO < 25% (resulting in 15/20 reported participants) and used the F4-A1 channel (Ying et al., 2025, Fig.S6) for consistency with Replication Dataset 1.

**Supplemental Table 2. The DTW analysis replication**

| Dataset | Main Dataset: Ultra-Long-Term Sleep project, Danish | Replication Dataset 1: Donders 2022, Dutch | | Replication Dataset 2: continuous multi-night sleep database, Chinese |
| --- | --- | --- | --- | --- |
| Device type | Subcutaneous EEG implant | Wearable EEG | Polysomnography | Polysomnography |
| Reference to the original study | Ahrens et al., 2024 | Jafarzadeh Esfahani et al., 2023 | | Ying et al., 2025 |
| No. participants | 20 | 27 | 24 | 15 |
| No. nights | 5656 | 322 | 68 | 45 |
| No. nights/participant | 282.8 | 11.9 | 2.8 | 3.0 |
| Channel(s) | TP9/TP10, T9/T10 and FT9/FT10 | latero-frontal | F3, F4 | F4-A3 |
| Dissimilarity (range) | 0.19 (0.16–0.24) | 0.28 (0.17–0.43) | 0.22 (0.12–0.38) | 0.28 (0.17–0.39) |
| Warping amount | 0.58 | 0.61 | 0.60 | 0.96 |
| Coefficient of variation | 0.35 | 0.40 | 0.28 | 0.36 |

*Dissimilarity was calculated by normalizing the DTW distances by the length of the time series (300 min). Between-subject range in dissimilarity is reported. CVs in dissimilarity across a year were calculated as the ratios between the standard deviation to the mean of dissimilarity between two times series within each participant. CV = 0.2 – 0.4 – medium variation, CV > 0.4 – high variation, CV – coefficient of variation.*

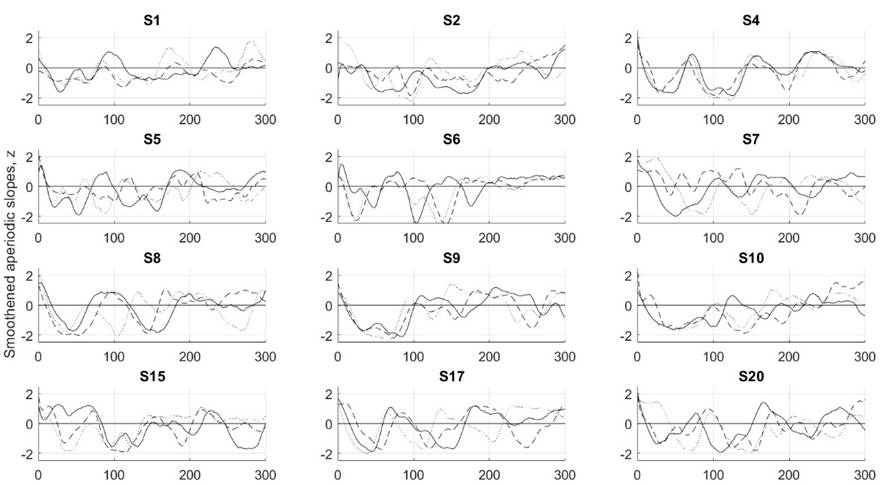

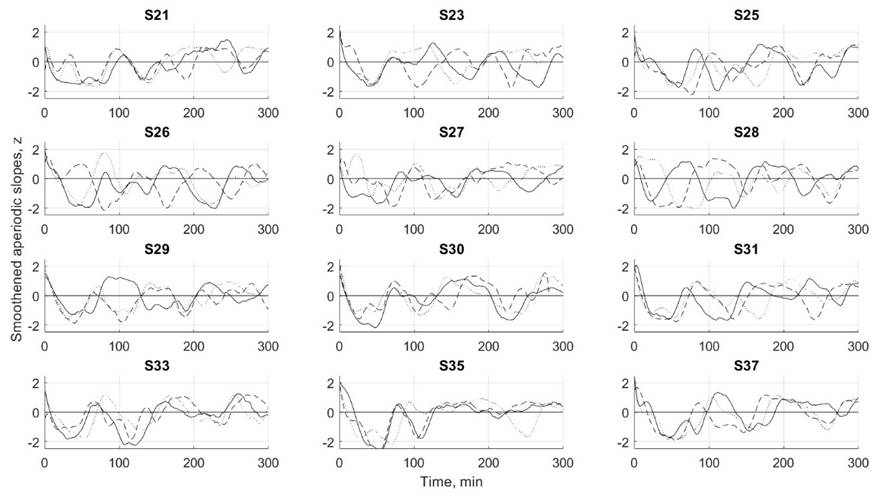

**Supplemental Figure 4. The macroscale unit of whole-night sleep. Polysomnographic Replication Dataset 1.** Each graph shows three 5h-long time series of z-normalized smoothened aperiodic slopes in one participant (see also Supplemental Table 2) averaged over F3 and F4 channels.

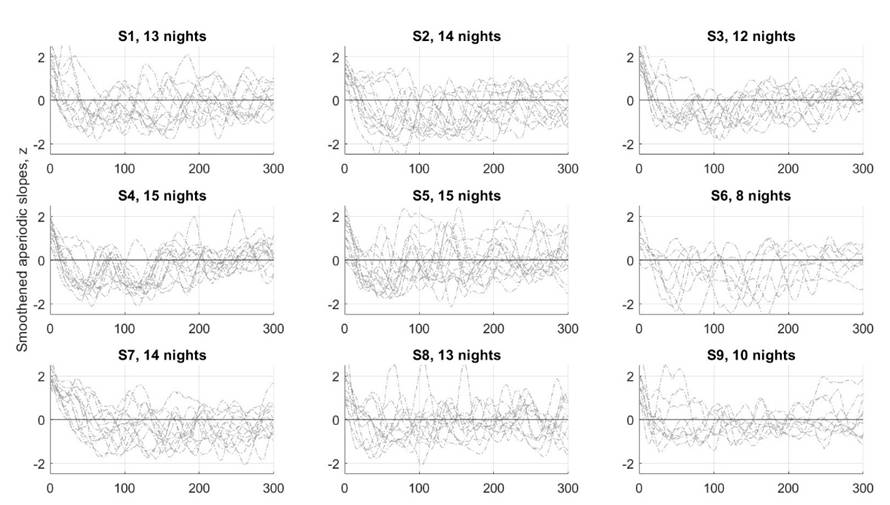

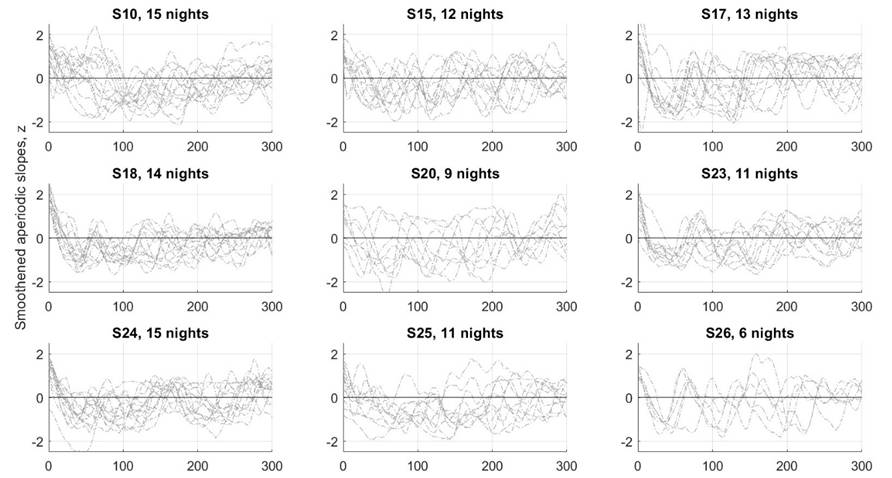

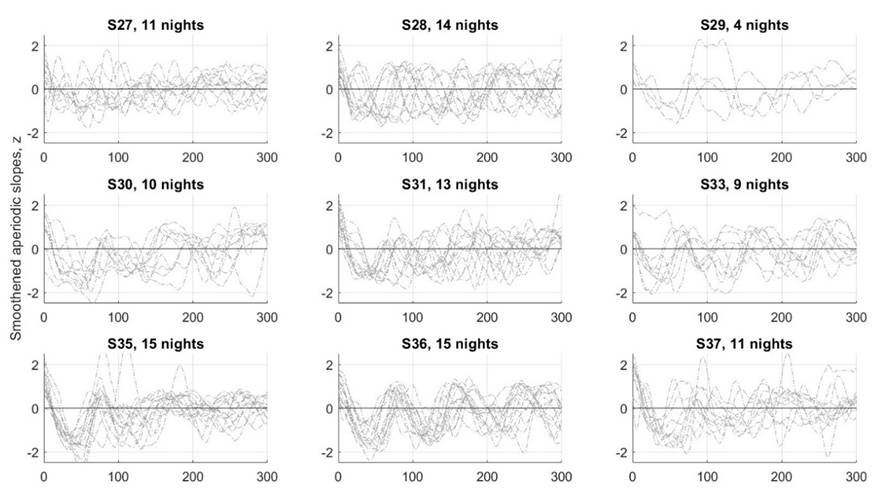

**Supplemental Figure 5. The macroscale unit of whole-night sleep. Wearable Replication Dataset 1.** Each graph shows up to fifteen 5h-long time series of z-normalized smoothened aperiodic slopes in one participant recorded with wearable device (Zmax, Hypnodyne, Sofia, Bulgaria) averaged over two latero-frontal channels and referenced to fronto-central channel (see also Supplemental Table 2).

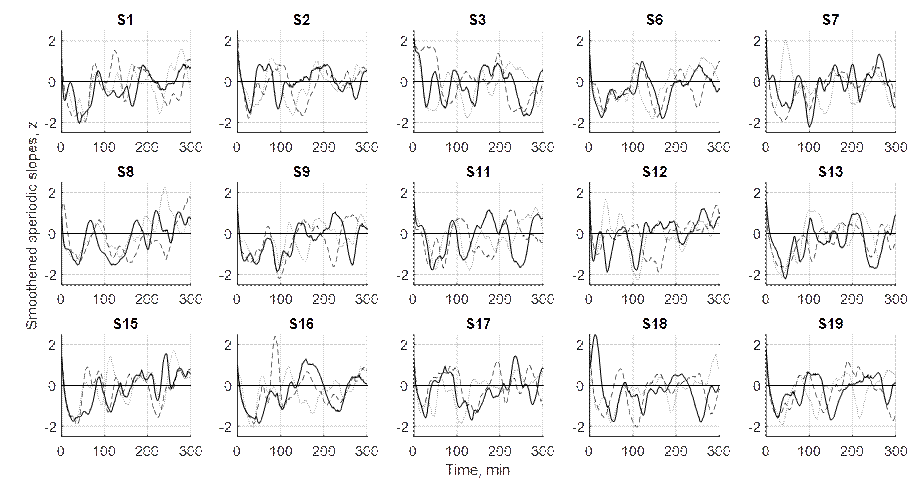

**Supplemental Figure 6. The macroscale unit of whole-night sleep. Polysomnographic Replication Dataset 2.** Each graph shows three 5h-long time series of z-normalized smoothened aperiodic slopes from the F4-A3 channel in one participant (see also Supplemental Table 2).
